# The antitumoral activity of TLR7 ligands is corrupted by the microenvironment of pancreatic tumors

**DOI:** 10.1101/2020.03.05.978767

**Authors:** Marie Rouanet, Hubert Lulka, Pierre Garcin, Martin Sramek, Delphine Pagan, Carine Valle, Emeline Sarot, Vera Pancaldi, Frédéric Lopez, Louis Buscail, Pierre Cordelier

## Abstract

Toll like receptors are key players in the innate immune system. Recent studies have suggested that they may impact the growth of pancreatic cancer, a disease with no cure. Among them, Toll like receptor-7 shows promise for therapy but may also promote tumor growth. Thus, we aimed to better understand the mechanism of action of Toll like receptor-7 ligands in pancreatic cancer, to open the door for clinical applications. *In vitro*, Toll like receptor-7 ligands strongly inhibit the proliferation and induce cell death by apoptosis of pancreatic cancer cells. *In vivo*, while Toll like receptor-7 agonists significantly delay the growth of aggressive tumors engrafted in immunodeficient mice, they instead surprisingly promote tumor growth and accelerate animal death in immunocompetent models. Molecular investigations revealed that Toll like receptor-7 agonists strongly increase the number of tumor-promoting macrophages to drive pancreatic tumorigenesis in immunocompetent mice. This is in stark contrast with Toll like receptor-7 ligands’ great potential to inhibit pancreatic cancer cell proliferation *in vitro* and tumor growth *in vivo* in immunosuppressed models. Collectively, our findings shine a light on the duality of action of Toll like receptor-7 agonists in experimental cancer models, and calls into question their use for pancreatic cancer therapy.

## Introduction

Pancreatic adenocarcinoma (PDAC) is a disease with no cure projected to become the second deadliest cancer worldwide by 2025^1^, with 5-year survival at less than 10%^2^. PDAC’s dismal prognosis is due to late diagnosis and to the lack of effective therapies^3^. Thus, there is an urgent need to define successful treatments to improve the prospects of patients diagnosed with PDAC.

Toll like receptors (TLRs) are pattern recognition receptors that are expressed on innate immune cells and in tumors^4^, including PDAC^5^. TLR ligands include bacterial and viral motifs (PAMPs for pathogen-associated molecular patterns), and damage-associated molecular patterns (DAMPs)^6^. TLRs recently emerged as promising targets, as they demonstrate therapeutic interest both on cancer cells and other cells in the tumor microenvironment (TME). Indeed, TLR activation can result in lethal autophagy, cell death by apoptosis or pyroptosis of cancer cells^7^. In addition, TLRs control tumor-promoting inflammatory signaling pathways common to both myeloid and lymphoid cells. Consequently, TLR manipulation has an impact not only on myeloid cells, but it also increases the activity, specificity and efficacy of adaptive immunity within the TME^7^. Recently, TLR2 agonists mimicking microbial signals were found to generate tumor-suppressive macrophages^8^. In another example, stimulating TLR7, an endosomal single strand (ss) RNA receptor associated with viral response and proinflammatory cytokines and type I interferon production^9^, inhibits programmed cell death protein 1 (PD-1) expression on T cells and promotes CD8- T-cell cytotoxic responses^10^. TLR7 agonists include benzapines and imidazolquinolines, with FDA-approved imiquimod and its more active version resiquimod (R848). Such small-molecule agonists demonstrated preliminary antitumoral efficacy but require further optimization to reach their full therapeutic potential^11^.

This is particularly true for PDAC, as TLR7 engagement demonstrates diametrically opposite effects on cancer cell lines and tumor outcome^12^. *In vitro*, TLR7 ligation results in both inhibition^13^ and stimulation of cancer cell proliferation^14^. *In vivo*, TLR7 promotes pancreatic carcinogenesis in mice by driving stromal inflammation^15^ and participates in microbiome-induced tumor promotion^5^. On the other hand, in interventional studies, TLR7 (adjuvant) therapy is often associated with beneficial immune antitumor response^16–18^. For instance, stromal rather than neoplastic TLR7 remodels tumor and host response to increase survival of mice with experimental PDAC, notably after improving cancer-associated cachexia^18^. On the contrary, Calore *et al.* recently demonstrated that TLR7 may also actively participate in cachexia, strongly arguing that interfering with TLR7 may serve as a potential therapy for the treatment of this highly debilitating syndrome associated to cancer^19^. Collectively, although TLR7 is expressed in cancer lesions^5^, mainly within the tumor stroma^18^, studies in well-defined experimental models are lacking to definitely address the therapeutic interest of TLR7 activation in PDAC management.

In this study, we investigated whether novel TLR agonists^20^ elicit anti-proliferative and anti-tumoral responses in immunodeficient or syngeneic orthotopic PDAC models. Remarkably, these models were generated with the same primary cell line, to determine TLR7 intrinsic action on neoplastic cells, but also on experimental tumors in the presence or not of a functional immune TME. We first demonstrate that *in vitro*, TLR7 agonists potently inhibit human and murine PDAC cell lines proliferation and induce cancer cell death by apoptosis. When tumors are engrafted in immunodeficient animals, TLR7 treatment translates into a significant delay in inhibition of experimental tumor progression and increase in mice survival. On the contrary, treating orthotopic PDAC tumors established in fully immunocompetent animals with TLR7 agonists results in stimulation of cancer cell proliferation, tumor promotion and decrease in mice survival. We found that tumors in immunocompetent mice treated with TLR7 agonists have a massive infiltration of macrophages expressing tumor-supportive markers. Collectively, our data demonstrate that TLR7 agonists have can be of interest to target PDAC cancer cells, but they also induce a tumor promoting TME, which strongly makes us question their potential as a therapy for patients with PDAC.

## Results

### TLR7 agonists inhibit PDAC cancer cells proliferation and induce cell death by apoptosis

For this study, we selected murine R211 cells derived from genetically engineered mouse models of PDAC with *KRAS* activating mutation and loss in *TP53*^21^. These cells are closely resembling tumor genetic and phenotypic landscapes and recapitulate tumor architecture and growth when implanted in mice pancreas. We found that human and mice pancreatic cancer cells express detectable levels of TLR2 and TLR7, with R211 expressing the highest levels of the candidate TLR (Supp. Figure 1A). Thus, R211 cells were treated with a panel of novel TLR2 and TLR7 agonists^20^. As positive controls, cells were left untreated, or treated with SSRNA40 (TLR7/8 agonist), and PAM3CSK4 (TLR2/1 agonist). We next performed non-invasive, longitudinal analysis of cell proliferation and induction of apoptosis using the Incucyte Zoom (Supp. Figure 1B-1E). We found that TLR7 agonists CL264 and ssRNA-40 only moderately inhibit PDAC cell proliferation (Fig. 1B and 1C, Table 1)). On the contrary, PAM3CSK4 (TLR2/1 agonist, Fig. 1A), CL307 and CL347 (TLR7 agonists, Fig. 1D and 1F), CL419 (TLR2 agonist, Fig. 1E) and CL553 (TLR2/7 agonist, Fig. 1G) impair PDAC cell lines cell proliferation (Table 1). However, CL347 demonstrates the strongest antiproliferative and proapoptotic activity (IC50=77μM±1.2μM, Table 1 and Fig. 2A). We further demonstrated CL347 proapoptotic effect by FACS analysis, as CL347 treatment significantly increases the number of apoptotic cells (Fig. 2B and 2C), and Western blot as CL347 induces a 2.3-fold increase in PARP cleavage (Fig. 2D). We confirmed CL347 antiproliferative effect by endpoint cell counting experiments in Mia PACA-2 (human PDAC), PKP16 (murine PDAC) and DT6606 (murine PDAC) cell lines (Supp. Figure 1F).

**FIGURE 1.**
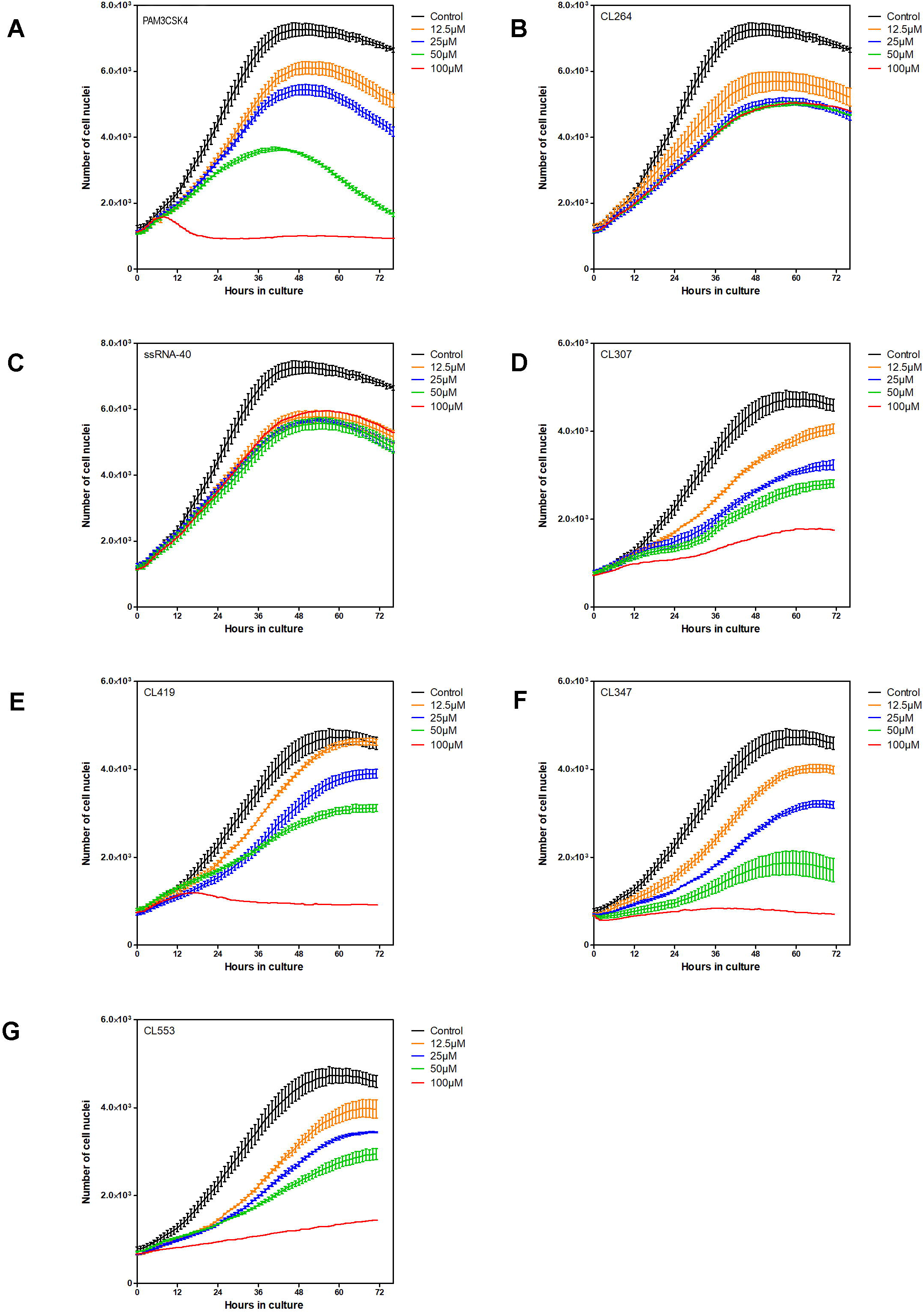

**Table 1:**
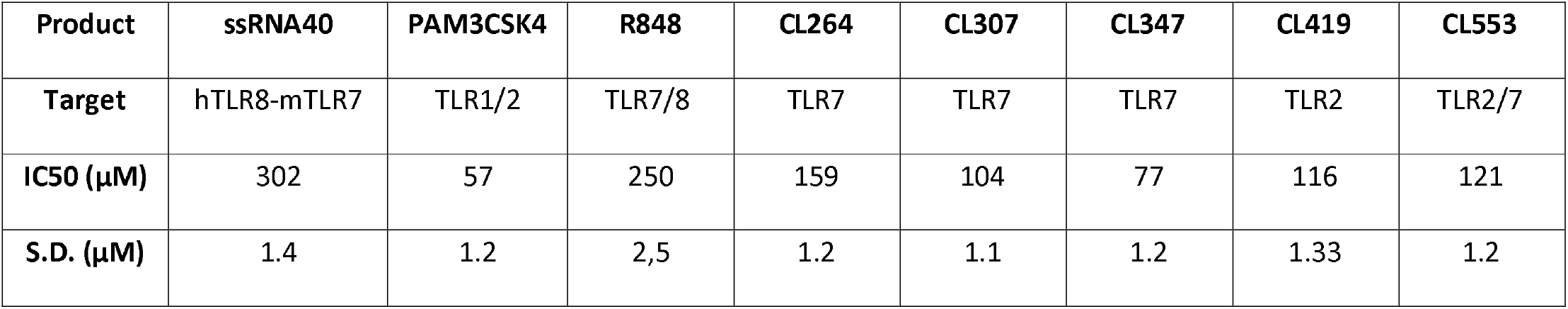
antiproliferative efficacy of TLR agonists on murine pancreatic cancer cell proliferation. Results are mean IC50 ±S.D. of 3 independent experiments.

| Product | ssRNA40 | PAM3CSK4 | R848 | CL264 | CL307 | CL347 | CL419 | CL553 |
| --- | --- | --- | --- | --- | --- | --- | --- | --- |
| Target | hTLR8-mTLR7 | TLR1/2 | TLR7/8 | TLR7 | TLR7 | TLR7 | TLR2 | TLR2/7 |
| IC50 (μM) | 302 | 57 | 250 | 159 | 104 | 77 | 116 | 121 |
| S.D. (μM) | 1.4 | 1.2 | 2,5 | 1.2 | 1.1 | 1.2 | 1.33 | 1.2 |

**FIGURE 2.**
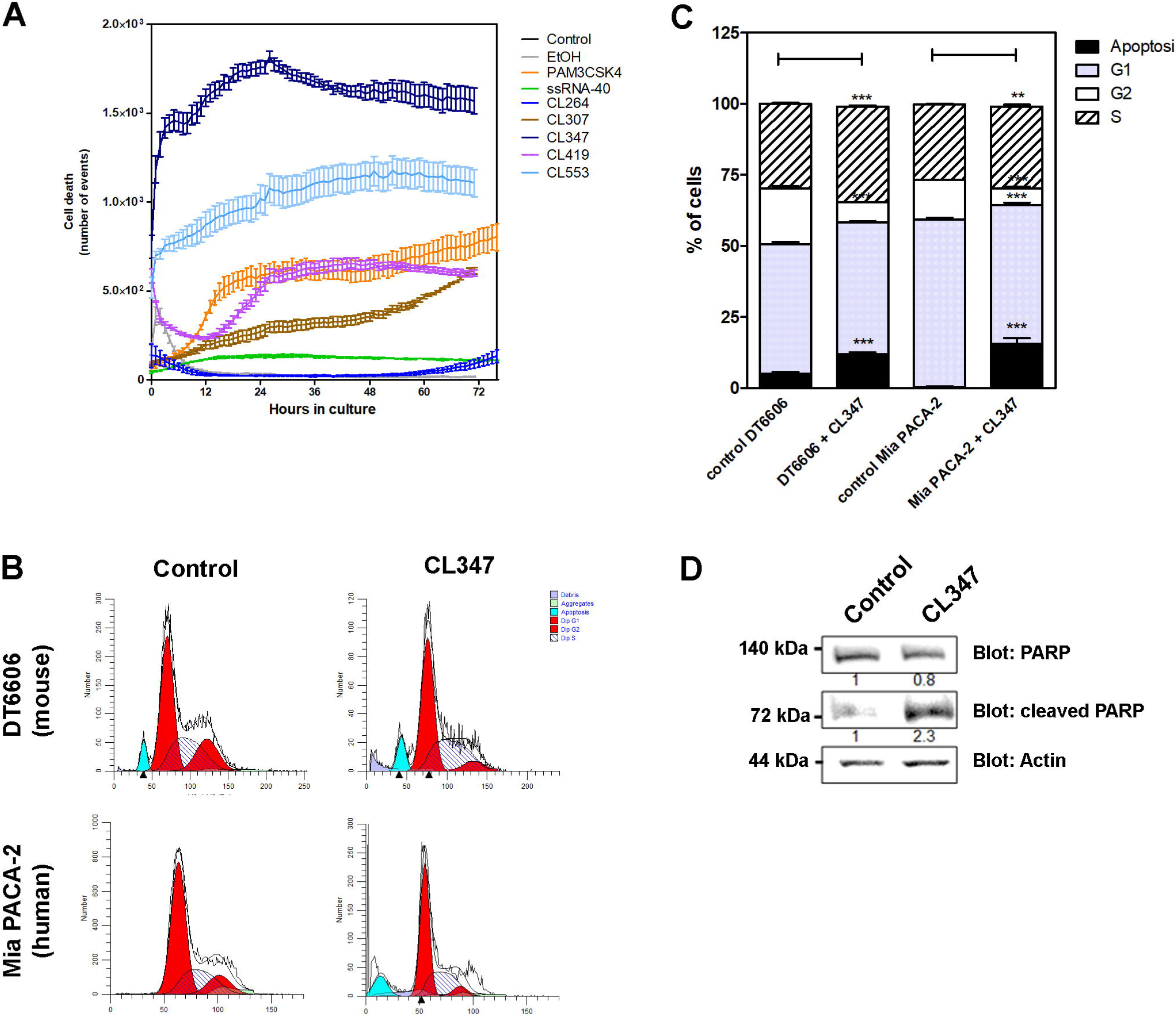

### *In vivo* TLR7 agonist administration inhibits PDAC cell proliferation and tumor progression and increases mice survival

The effect of TLR7 agonists on tumor progression is multifaceted, with an impact on both neoplastic cells and the TME. We first addressed the cell intrinsic activity of the TLR7 agonist CL347 on PDAC models, in the absence of the immune microenvironment. NOD scid gamma (NSG) mice were implanted with CL347-responsive R211 cells expressing red-shifted, firefly luciferase (R211-RSLucF). Tumor growth was monitored non-invasively by detecting luciferase bioluminescence using the Ivis Spectrum. Animals with exponentially growing tumors were treated intraperitoneally with different doses of CL347, and subsequently characterized by analyzing tumor growth and survival. By 15 days, all mice treated with placebo developed detectable tumors (r.l.u. > 1×10e10, Fig 1A and 1F). Treatment of orthotopic PDAC tumors with CL347 resulted in a dose-dependent delay in tumor growth as compared to placebo (Fig.3B-D). Importantly, mice treated with the highest dose of CL347 (80μg) had no detectable bioluminescence up to 25 days following treatment (Fig. 1D and 1F). In addition, we found that 21 out of 22 mice (95%) receiving CL347 survived during the experiment while only 15% of mice from the placebo group survived (Fig.1E). Interestingly, 75% of mice treated with the highest dose of TLR7 agonist were free of ascites (data not shown). At the end of the experiment, mice were killed, and tumors were sampled. Immunohistochemistry detection of Ki67 demonstrated that CL347 treatment significantly reduces the number of proliferating cancer cells (Fig. 1G and 1H, p<0.005). Importantly, significant concerns exist about TLR agonists toxicity when administered systemically, despite beneficial effects on tumor response. However, we did not observe mice weight loss following treatment with CL347 during this study (data not shown). Collectively, these results demonstrate the antiproliferative action of TLR7 agonists *in vivo*, which translates into tumor growth control and improved survival in experimental immunodeficient models of PDAC.

**FIGURE 3.**
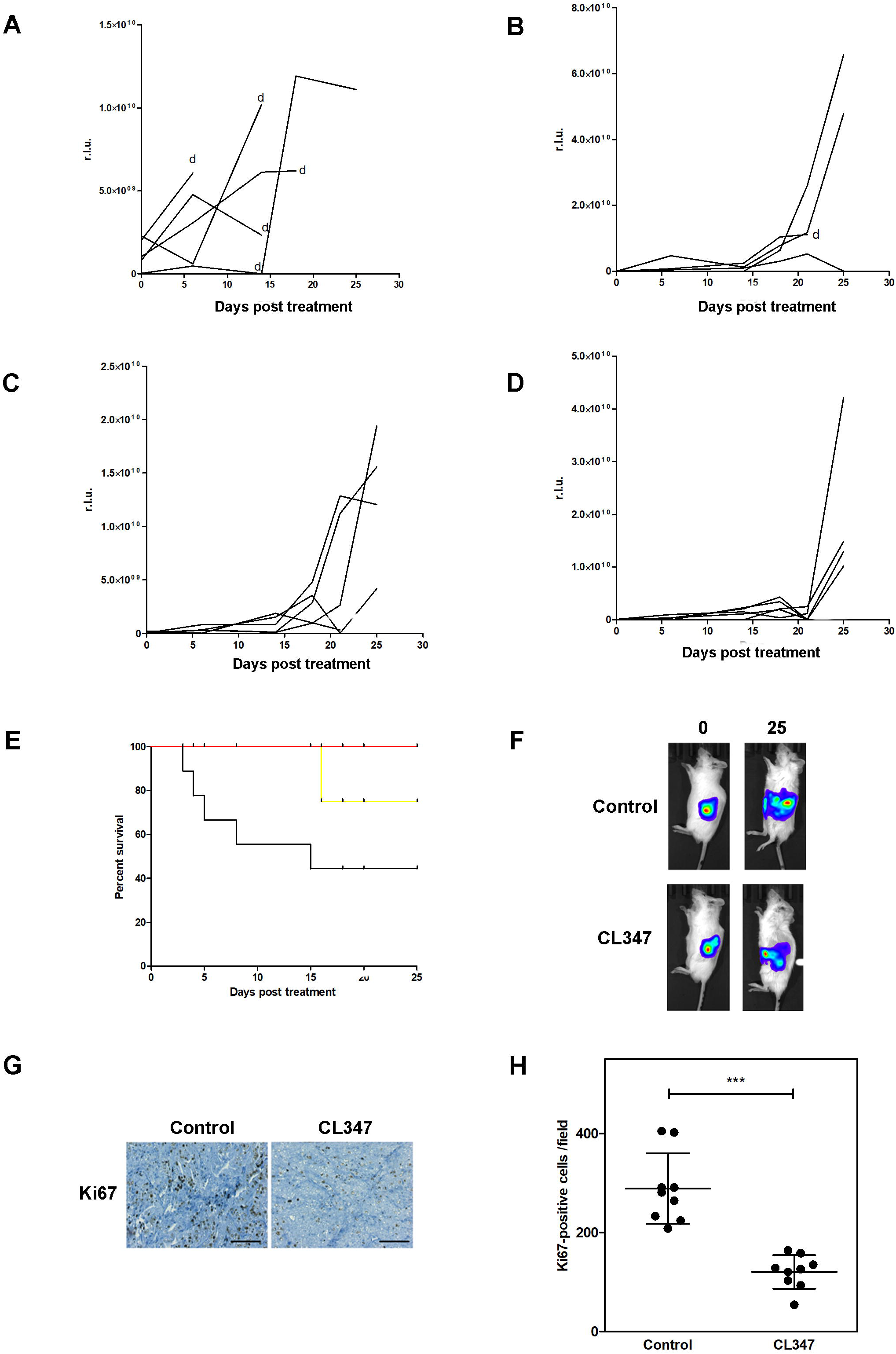

### TLR7 treatment favors tumor progression in immunocompetent hosts

TLR agonists are used against a variety of malignancies to induce antitumoral immunity^7^. We explored whether TLR7 agonist CL347 could activate tolerant host immune system to eradicate experimental PDAC. To this end, R211-RSLucF cells were engrafted in the pancreas of immune competent mice and tumor growth was monitored non-invasively as described before. As expected, host response significantly delayed the increase in tumor burden, suggesting tumor control by immune cancer killing cells (Fig. 4A). To our surprise, we found that TLR7 treatment dramatically accelerates tumor progression (Fig.4B-D, Fig. 4F), and shortens mice survival (Fig. 4E), in a dose-dependent manner. By immunohistochemistry, we found that TLR7 treatment leads to tumors with more aggressive phenotypes and higher proliferative index (Fig. G-H). Considering the well-described effect of TLR7 on immune cell populations, R211 tumors were analyzed for lymphocytic (CD3) and myeloid (F4/80) markers. On one hand, we found that CL347 treatment significantly decreases the number of lymphocytes infiltrating R211 tumors (Fig. 5A and 5B); on the other hand, CL347 massively increases the number of F4/80 positive cells in tumors (Fig. 5A and 5C). Analysis of differential expression in tumors treated with CL347 was performed using Nanostring mouse inflammation panel and revealed the presence of signatures characteristic of abnormal innate inflammatory response (p value = 3.57×10^−16^, Bonferrroni correction), and overrepresentation of the complement pathway (p value = 3.153×10^−12^, Bonferrroni correction) (Supplemental file 1). The most striking changes in gene expression involve downregulation of the expression of cytokines of different categories:(i) cytokines regulating lymphocyte trafficking (MCP-1/CCL2, logFC= −1.96, p val. = 3.3E-07), (ii) cytokines produced by tumoricide, M1 tumor-associated macrophages (CXCL10,log FC= −1.93, p val.= 2.8E-05), (iii) cytokines involved in neutrophil activation (CXCL5, logFC= −1.8, pval= 4.6E-04) and eosinophils attraction (CCL24, logFC=−1.66, pval=1E-03, Fig.4D and 4E). On the other hand, type II, protumoral tumor associated macrophage markers (CD163 log FC= 2.9, p val.= 1.5E-11, MASP1 log FC= 3.6, p val.= 2.8E-09 and CHI3L3 log FC= 5.01, p val.= 6.1E-6) are significantly induced in tumors upon treatment by CL347 (Fig. 4D and 4E).

**FIGURE 4.**
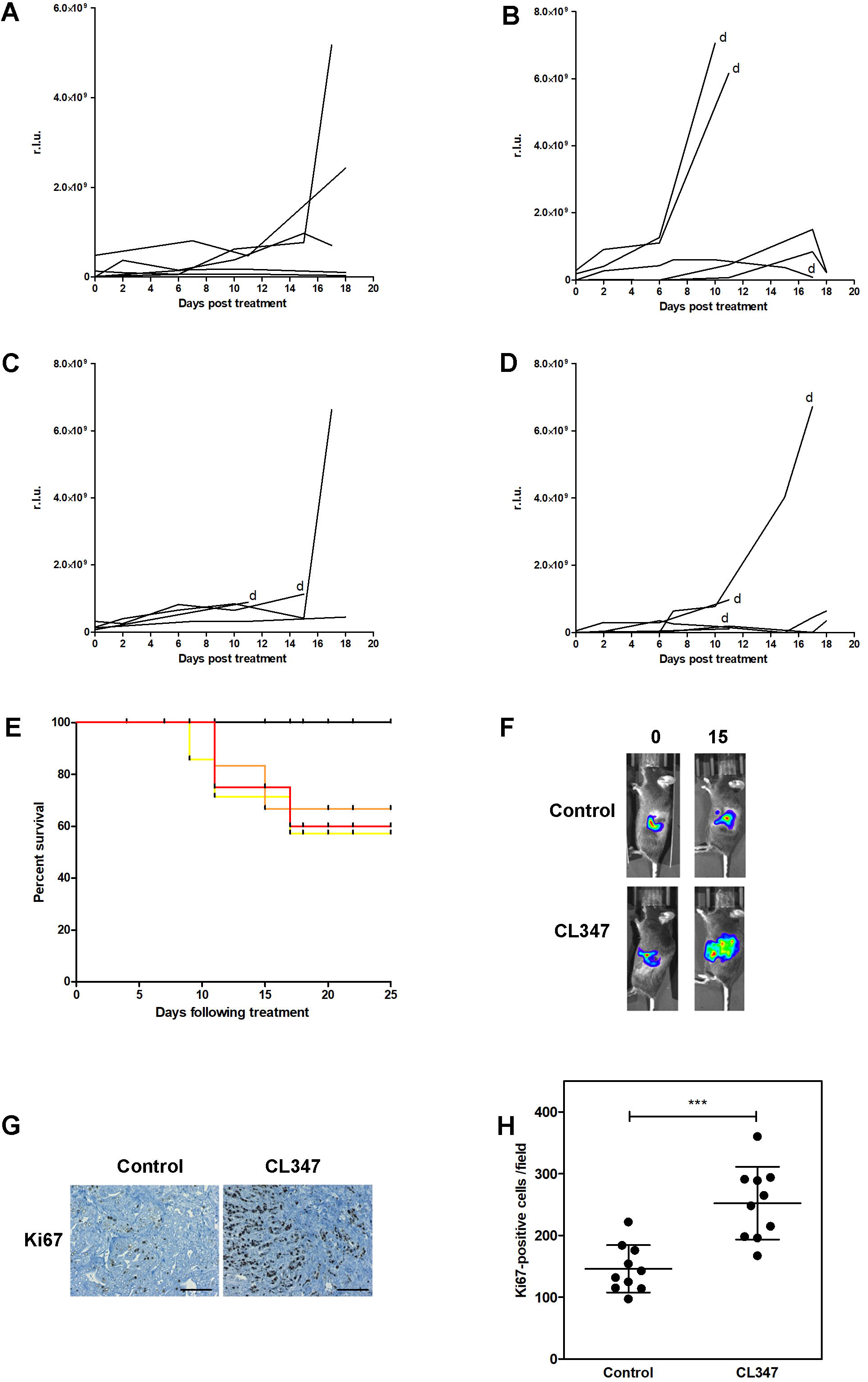

**FIGURE 5.**
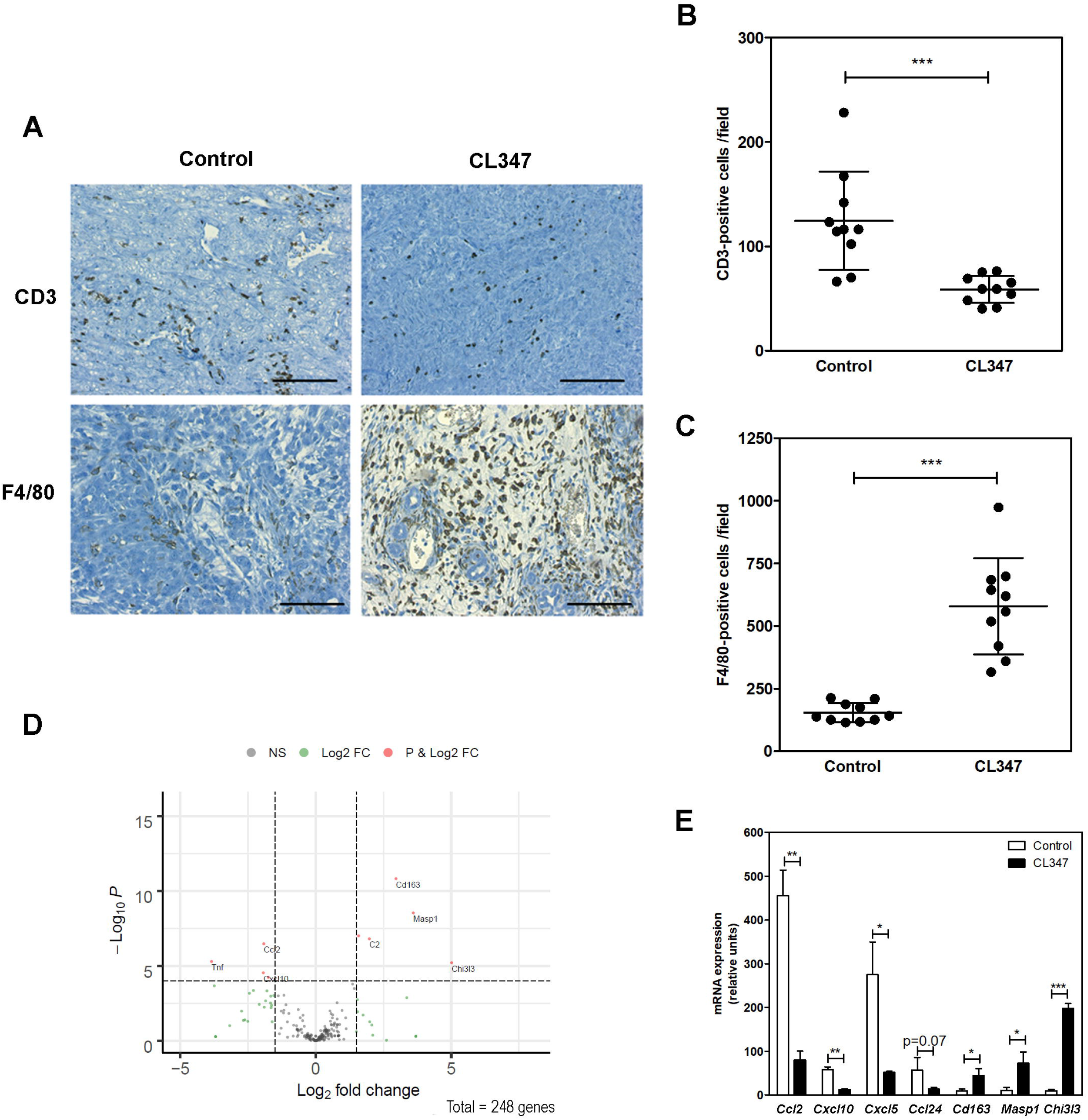

Collectively, our data demonstrate that TLR7 ligation inhibits PDAC cell proliferation *in vitro* and *in vivo* in immunodeficient models of PDAC. However, in immunocompetent experimental models closely resembling human disease, TLR7 activation is protumoral and shortens mice survival. Molecular explorations point to a crucial role of type-II macrophages in TLR7 protumoral activity.

## Discussion

PDAC is one of the few deadly diseases that has not yet been defeated by immunotherapies^3^. In this context, TLR ligands are of high therapeutic interest as they show promising antitumoral activity either by directly inhibiting PDAC cell proliferation or inducing innate immune response against the tumor^12^.

Unfortunately, multiple lines of evidence point to a more complex picture, with recent studies reporting opposite outcomes following treatment of experimental PDAC with TLR agonists. In this study, we confirm that TLR7 agonists have intrinsic antiproliferative and pro-apoptotic activities in cell lines, which ultimately translate into tumor growth control in immunodeficient mice. However, we also highlight the Janus face of TLR7 on tumor growth as acute TLR7 ligands administration remodels the immune microenvironment of PDAC experimental tumors ultimately increasing tumor burden and shortening mice survival. We found that administration of TLR7 agonists decreases intratumoral T-cell infiltration and promotes type-2 macrophage colonization of tumors. Based on these data, we report that despite a valuable inhibitory action on neoplastic cells, acute treatment with TLR7 agonists also promotes tumor supportive host immune response. We also provide the starkest possible reminder that PDAC tumors cannot be considered as an assembly of neoplastic cells, given the crucial importance of stromal reactions associated with this tumor type^2^.

Our study is consistent with recent works from the literature illustrating the duality of action of TLR7 in PDAC models. As stated before, PDAC is an aggressive cancer that interacts with stromal cells to produce a highly inflammatory TME that promotes tumor growth and invasiveness. In a immunocompetent mouse model of PDAC, genetic ablation of TLR7 within inflammatory cells protects from neoplasia^15^. In a sibling study, Michaelis *et al.* remarkably demonstrated that host rather than neoplastic TLR7 is necessary for TLR agonists’ beneficial antitumoral effect, notably after engagement of an antitumoral immune response^18^. These authors also raise substantial awareness after showing that chronic administration of the TLR7 ligand R848 could also increase tumor growth^18^. Interestingly, these authors also performed short-term, so called “burst” experiments with TLR7 ligands, a situation that more closely resembles our “acute” regimen. They found that survival was only modestly increased as compared to placebo. Unfortunately, the tumor immune microenvironment was not characterized in the latter settings, so questions regarding massive protumoral immune response following acute TLR7 treatment are still open. Collectively, these results suggest that the dose and the frequency of TLR7 administration may also have different consequences on experimental PDAC tumor growth outcome.

Further studies are needed to better understand the molecular mechanisms behind TLR7-driven establishment of this protumoral immune microenvironment. In mouse, TLR7 is predominantly expressed in plasmacytoid dendritic cells and B cells, with context dependent expression in a subset of macrophages and T cells^22^. Our results strongly suggest that TLR7 treatment populates PDAC tumors with tumor-associated macrophages (TAMs) with pro-tumor state, a situation at the antipodes of ideal scenarios for immunotherapy. Indeed, these cells act as a powerhouse for tumor angiogenesis and metastasis, by displaying an immunosuppressive (M2) phenotype and secreting abundant pro-tumor cytokines^23^. Among TLR family members, TLR2 activation of peripheral monocytes triggers differentiation into DC-SIGN+ macrophages, with robust M1 anti-tumoral phenotype^24^. Along the same line, recent work reports that acGM-1.8, a specific TLR-2 agonist, generates macrophages with strong anti-tumor potential in mice^8^. For TLR7, a recent report shows that R848, an agonist of TLR7 and TLR8, is a potent driver of the M1 phenotype *in vitro* and that R848-loaded nanoparticles target TAMs *in vivo* to impede colon tumor and melanoma growth in mice^25^. In our hands, R848 demonstrates poor antiproliferative activity, *in vitro* (data not shown), but unfortunately, PDAC models were not investigated during the aforementioned study. It is tempting to speculate that M2 macrophages could be triggered/attracted by apoptosis in the tumor induced by CL347. In this respect, the milder effect of TLR7 ligand CL307 on cell proliferation and apoptosis induction we identified during this study could be more beneficial. Finally, it would be of interest to determine the origin of TAMs following administration of TLR7 in animals bearing PDAC tumors. Indeed, one simple explanation supported by our result would be that TLR7 treatment favors inflammatory monocytes conversion into M2 TAMs. On the other hand, a recent report nicely demonstrates that tissue-resident macrophages are an essential source of TAMs in murine PDAC models^26^. Such embryonically derived TAMs exhibit a pro-fibrotic transcriptional profile, indicative of their role in producing and remodeling molecules in the extracellular matrix^26^. Interestingly, we found that the expression of several metalloproteinases is induced following CL347 treatment of immune competent PDAC tumors, despite the lack of statistical significance in this trend.

Taken together, our results highlight the importance of evaluating candidate molecules using animal models that faithfully recapitulate tumors in patients, as therapeutic outcome must be conceived of as the sum of effects on the cancer cells and the host (TME) in which the disease takes place. Here, we shine a light on the duality of action of TLR7 agonists in experimental PDAC models, and call into question the use of TLR7 agonists for PDAC therapy.

## Materials and methods

### Reagents

PAM3CSK4 (TLR1/2 agonist), SSRNA-40 (TLR7/8 agonist), CL264 (TLR7 agonist), CL307 (TLR7 agonist), CL347 (TLR7 agonist), CL419 (TLR2 agonist) and CL553 (TLR2/7 agonist) were purchased from Invivogen (Toulouse, France). A complete description of the different compounds’ affinity and efficacy is available at www.invivogen.com. Lentiviral plasmids encoding for Red-shifted luciferase (RediFect Red-FLuc, RSLucF) or for NucLight Green (NucG) were from Perkin Elmer (Waltham, MA, USA) and Sartorius (Essen Biosciences, Hertfordshire, UK), respectively. Antibodies used in this study were from Cell Signaling Technology (total PARP, ref. 9532, cleaved PARP ref. 5625, b-actin ref. 8H10D10) and used following the manufacturer’s recommendation.

### Cell culture

All murine pancreatic cancer cells are kind gifts of D. Sauer (TUM, Munich, Ger). R211 cells derive from spontaneous tumors generated in the KPC mice model (LSL-KrasG12D/+; LSL-Trp53R172H/+; Pdx-1-Cre). DT6606 cells derive from spontaneous tumors generated in the KC mice model (LSL-KrasG12D/+; Pdx-1-Cre). PKP16 cells derive from spontaneous tumors generated in the KP16C mice model (LSL-KrasG12D/+; LSL-Tp16/INK4A/+; Pdx-1-Cre). Mia PACA-2 (ATCC CRL-1420) cells were purchased from LGC standards (UK). Cells were grown in DMEM 4.5g/L glucose medium supplemented with 10% fetal calf serum (FCS), L-glutamine, antibiotics (Life Technologies) and plasmocyn (Invivogen, complete medium) at 37°C in humid atmosphere with 5% CO2. Cell cultures were certified mycoplasma-free (PlasmoTest kit, Invivogen).

### Lentiviral vector production and cell transduction

All replication defective, self-inactivating lentiviral vectors were generated in a BSL-3 facility (Vectorology platform, INSERM U1037, Toulouse, France) as previously described ^1^. Briefly, transient transfection of HEK-293FT cells with packaging and lentiviral vector plasmids were performed using LENTI-Smart INT kit (InvivoGen) following manufacturer’s recommendations. All batches were verified replicative virus-free. Lentiviral vector concentrations were quantified by p24 ELISA (Innotest, Ingen, Paris). Cells were seeded at a density of 10^4^ cells per well in a 48 well-dish. After 24h, cells were incubated with 150 ng of p24-equivalent of lentiviral vectors in the presence of protamine sulfate (4 μg/mL) for 16h. Transduced cells were selected for 3 weeks using puromycin (5 μg/mL,-InvivoGen).

### Cell proliferation

IncuCyte ZOOM Live-Cell Imaging system (Sartorius) was used for kinetic monitoring of cell proliferation and cytotoxicity of TLR agonists in murine pancreatic cancer cells. Briefly, R211-NucG cells were seeded at 10^4^/well in 96-well black-walled plates. Cells were treated with increasing concentrations (12.5-100 μM) of TLR agonists in complete medium in the presence of IncuCyte® Cytotox Red (Sartorius), following the manufacturer’s recommendation. R211 nucleus is labelled in green, and IncuCyte® Cytotox Red labels dead cells yielding red fluorescence. The plate was scanned and fluorescent and phase-contrast images were acquired in real time every hour from 0 to 72 hours post treatment. Normalized Green and Red Object Count per well at each time point were generated by IncuCyte ZOOM software and plotted using GraphPad Prism v8.3.1 (549) (Graphpad Software, San Diego, CA, USA). For cell counting studies, DT6606, PKP16 and Mia PACA-2 cells were seeded in triplicate at a concentration of 5.10^4^/well of 6 well-plate. After 24h (t=0), cells were counted using a Z1 Coulter® Particle Counter (Beckman Coulter™), then treated with TLR ligands at the indicated dose. Cells were subsequently counted after 72h in culture.

### Western blotting

Cell pellets were incubated in RIPA buffer supplemented with 10 μL/mL protease inhibitor (Sigma-Aldrich). After 15 min on ice, samples are centrifuged (15 min-12 000rpm, 4°C). Protein fractions were denaturated in Laemmli buffer after heating at 95°C for 5 min. Proteins were separated by SDS-PAGE (Sodium Dodecyl Sulfate Polyacrylamide Gel Electrophoresis) and transferred onto nitrocellulose membrane (Bio-Rad) using TransBlot Turbo (Bio-Rad) apparatus. After membrane saturation and primary/secondary antibody incubation, protein expression was detected using ClarityTM Western ECL Substrate (Bio-Rad) and Chemi-DocTM XRS+ (Bio-Rad) apparatus. Signal intensities were quantified using Image Lab (Bio-Rad) software.

### Flow cytometry

For cell cycle analyses, cells were fixed in 70% ethanol during the exponential growth. Fixed cells were treated with RNAse A (10 μg/mL) and propidium iodide (20 μg/mL) (Sigma-Aldrich) for 15 min at 37°C. Data were acquired using the MACS Quant® VYB cytometer (Miltenyi Biotech) and analyzed with MACS Quant and ModFit software.

### Gene expression analysis

For Quantitative reverse transcription PCR studies (RT-qPCR), total RNA was isolated cell lines with TRIzol® Reagent (Life Technologies) according to supplier’s instructions. One to five μg of total RNA were reverse transcribed into cDNA using RevertAid H minus Reverse Transcriptase kit (Thermo-Scientific) according to manufacturer’s recommendations. Duplicate RT-qPCR assays were carried out in a StepOnePlus™ Real-Time PCR System (Applied Biosystems) with SsoFast™ EvaGreen® supermix (Bio-Rad) and specific primers (see primers listed in Supplemental Materials and Methods 2). Relative quantity of mRNA was calculated by the comparative threshold cycle (CT) method as 2^−ΔCT^, where ΔCT = CT *candidate gene* mRNA – CT *Reference* mRNA. *β*-Actin was used for normalization. Primers for TLR2 and TL7 expression studies are the following: human TLR2-Forward: TTG ATG ACT CTA CCA GAT GCC, human TLR2-Reverse: CAA ATG AAG TTA TTG CCA CCA G, human TLR7-Forward: TCT GAC TAA CCT GAT TCT GTT CTC, human TLR7-Reverse: GTG ATA TTA GAC GCT GAT ACC C, murine TLR2-Forward: GTC TTT CAC CTC TAT TCC CTC C, murine TLR2-Reverse: CCC TCT ATT GTA TTG ATT CTG CTG, murine TLR7-Forward: CGG TGA TAA CAG ATA CTT GGA C, murine TLR7-Reverse: AGA GAT TCT TTA GAT TTG GCG G.

For Nanostring quantification of gene expression, tumors were sampled and minced in liquid nitrogen. Total RNA was extracted with TRIzol® Reagent (Life Technologies) according to supplier’s instructions. One hundred nanograms of RNA were analyzed using the murine inflammation panel according to supplier’s instructions. Genes with reads below 50 were not further considered. Raw data from Nanostring was performed using the NanoStringDiff package (NanoStringDiff_1.14.0 Biobase_2.44.0). Differential Gene Expression analysis was conducted using the package (see process_nanostringFinal.R). Volcano plots were generated with EnhancedVolcano package (EnhancedVolcano_1.2.0). We performed enrichment analysis using ToppGene, accessed on 19th July 2019^2^.

### *In vivo* experiments

Experimental procedures performed on mice were approved by the ethical committee of INSERM CREFRE US006 animal facility and authorized by the French Ministry of Research: APAFIS#3600-2015121608386111v3. R211-RSLucF cells (5.10^4^ cells) were injected into the pancreas of NOD scid gamma (NSG) or C57bl6 mice (n=6 per group). Tumor growth monitoring was performed non-invasively twice a week by injecting Xenolight D-Luciferin (Perkin Elmer) and recording relative light units (r.l.u.) using the Ivis Spectrum (Perkin Elmer), following the manufacturer’s recommendations. Approximately two weeks following tumor induction, mice with exponentially growing tumors were injected intraperitoneally with 100μl of placebo (PBS with 5% glucose) or CL347 diluted in PBS with 5% glucose at the indicate dose (20μg, 40μg and 80μg). Tumor growth was monitored non-invasively as mentioned before at the indicated time point. Mice were killed 25 days (immunodeficient model) or 15 days (immunocompetent model) following treatment.

### Immunohistochemistry

Tumors were harvested and fixed in formalin. Four-micrometer-thick sections were prepared from paraffin-embedded sections and rehydrated. Ki67 staining was performed as previously described^3^. For F4/80 and CD3 staining, antigen retrieval was performed using Proteinase K (DakoCytomation) according to manufacturer’s recommendations or using Tris-EDTA, respectively. Slides were incubated one hour at room temperature with F4/80 antibodies (Thermo Scientific, ref. MA1-91124) diluted 1:100 or CD3 antibodies (Spring biosciences, Clone SP7, ref. M3070) diluted 1:200 in Antibody diluent solution (DakoCytomation). After several washes in PBS, secondary, goat anti-rat (mouse adsorbed, STAR131b) or goat anti-rabbit-HRP, both diluted 1:50 in Antibody diluent solution were applied for thirty minutes at room temperature, for F4/80 or CD3 staining, respectively, followed by horseradish peroxidase streptavidin (dilution 1:500, SA-5004, Vector). Slides were quickly washed twice in PBS and incubated in AEC+ reagent and counterstained with Mayer's hematoxylin (DakoCytomation). After washing in PBS, slides were mounted with Vectashield (Vector). Immunostaining was recorded with an AXIO optical microscope (Zeiss) equiped with a colour AXIOCAM camera 105 (Zeiss) and quantified using ImageJ.

### Statistical analysis

Data were analysed by 2-tailed, unpaired Student’s t-test using a multiple statistics Graph Pad Prism 8 software package and a difference was considered significant when p value was lower than 0.05. Mean values are given ± SD. Number of independent experiments is indicated in the figure legends. *, ** and *** indicate a p value < 0.05, <0.01 and <0.001, respectively. For Nanostring data analysis, we analysed first the genes that were highest in logFC (top30) irrespective of P-val and then gene that were Up and had p-val <0.1.

## Acknowledgments

This work was supported by grants from Region Occitanie (grant number 13053110 and 15052181) and CHU Toulouse (M.R.). The authors would like to thank Dr. M. Dufresne for immunohistochemistry protocols and Drs. F. Vernejoul and J. Torrisani for helpful discussions.

## Author Contributions

Conceptualization, P.C. and L.B.; investigations M.R., P.G., H.L., E.S., C.V., and M.S.; supervision, P.C.; writing – original draft, P.C.; writing – review & editing F.L., M.S. and V.P.; funding acquisition, P.C.; data analysis, L.B., P.C., V.P. and F.L.

