## Supplementary material for "The antitumoral activity of TLR7 ligands is corrupted by the microenvironment of pancreatic tumors": Figure legends

**Figure 1. Antiproliferative activity of TLR ligands on pancreatic cancer cells.** R211 cells derived from spontaneous murine pancreatic tumors and genetically engineered to express fluorescent nuclear reporter (R211-NucGreen) were seeded in 96-well plates and treated with PAMS3CSK4 (**A**), CL264 (**B**), SSRNA-40 (**C**), CL307 (**D**), CL419 (**E**), CL347 (**F**) and CL553 (**G**) at the indicated dose. Control cells were treated with medium only or TLR agonist solvent (water, ethanol). Cells nuclei were imaged and numbered non-invasively every hour for three days using the Incucyte Zoom as described in Materials and Methods. Results are mean ± SD of triplicates and representative of three independent experiments.

**Figure 2. TLR7 ligand CL347 induces massive cell death of pancreatic cancer cell lines. A.** R211-NucGreen cells derived from spontaneous murine pancreatic tumors were seeded in 96-well plates and treated with 100µm of PAMS3CSK4, CL264, SSRNA-40, CL307, CL419, CL347 and CL553. Control cells were treated with medium only or TLR agonist solvent (water, ethanol). Cells were incubated with cytotox red reagent for counting dead cells non-invasively every hour for three days using the Incucyte Zoom as described in Materials and Methods. Results are mean ± SD of triplicates and representative of three independent experiments. DT6606 cells derived from spontaneous murine pancreatic tumors (top) and Mia PACA-2 cells, derived from human pancreatic tumors were seeded in 60-mm dishes and treated for 48 hours with 50µM of CL347 and analyzed for cell cycle phase distribution by fluorescence-activated cell sorting. **B**. representative of three independent experiments. **C**. Results are mean ± SD of three independent experiments. **: p<0.01, ***: p<0.005. **D**. R211-NucGreen cells derived from spontaneous murine pancreatic tumors were seeded in 100-mm dishes and treated with 50µm of CL347 for forty-eight hours. Cells were lysed and soluble proteins were analyzed by Western blot for total Poly (ADP-ribose) polymerase (PARP) PARP-1 protein (top), cleaved PARP-1 (middle). Actin was used as loading control (bottom). Fold increase in PARP expression and cleavage calculated using actin as a control is indicated. Representative of three independent experiments.

**Figure 3. Antitumoral activity of CL347 TLR7 ligands in immunodeficient models of pancreatic cancer**. R211 cells derived from spontaneous murine pancreatic tumors and genetically engineered to express Red-shifted luciferase reporter gene (R211-RSLuc) were engrafted in the pancreas of athymic (NOD scid gamma) mice as described in Materials and Methods. Fifteen days following engraftment, mice received PBS placebo (**A**) or a total of 20µg (**B**), 40µg (**C**) and 80µg (**D**) of CL347 in 100µl of 5% glucose intraperitoneally. As control, mice were injected with 100µl of 5% glucose. Six mice were used per group. Tumor growth was monitored non-invasively using the Ivis spectrum and quantified as relative light units (r.l.u.) using Living image software. Results are representative of two independent experiments. d stands for death and indicates that mice had to be euthanized due to severe conditions related to tumor growth. **E**. Survival rate in control mice (black) and mice treated with 20µg (yellow), 40µg (orange) and 80µg (red) of CL347. **F**. Representative luminescence in mice treated or not by 80µg of CL347, before and twenty-five days following treatment. At the end of the experiment, control mice and mice treated by 80µg of CL347 were killed, tumors were sampled and analyzed for Ki67 proliferation marker. **G**. Representative Ki67 staining in control and mice treated by 80µg of CL347, 25 days following treatment. Scale bar: 100µm. **H**. Quantification of Ki-67 positive cells per field in control and mice treated by 80µg of CL347. Results are mean ± S.D. of ten fields from three different tumors. ***: p<0.005.

**Figure 4. TLR7 ligands treatment accelerate pancreatic carcinogenesis in immunocompetent mice.** R211 cells derived from spontaneous murine pancreatic tumors and genetically engineered to express Red-shifted luciferase reporter gene (R211-RSLuc) were engrafted in the pancreas of C57/BL6 mice as described in Material and Methods. Fifteen days following engraftment, mice received PBS placebo (**A**) or a total of 20µg (**B**), 40µg (**C**) and 80µg (**D**) of CL347 in 100µl of 5% glucose were injected intraperitoneally. As control, mice were injected with 100µl of 5% glucose. Six mice were used per group. Tumor growth was monitored non-invasively using the Ivis spectrum and quantified as relative light units (r.l.u.) using Living image software. Results are representative of two independent experiments. d stands for death and indicates that mice had to be euthanized due to severe conditions related to tumor growth. **E**. Survival rate in control mice (black) and mice treated with 20µg (yellow), 40µg (orange) and 80µg (red) of CL347. **F**. Representative luminescence in mice treated or not by 80µg of CL347, before and fifteen days following treatment. Fifteen days following treatment, control mice and mice treated by 80µg of CL347 were killed, tumors were sampled and analyzed for Ki67 proliferation marker. **G**. Representative Ki67 staining in control and mice treated by 80µg of CL347. Scale bar: 100µm. **H**. Quantification of Ki-67 positive cells per field in control and mice treated by 80µg of CL347. Results are mean ± S.D. of ten fields from three different tumors. ***: p<0.005.

**Figure 5. TLR7 ligands increase the number of type II macrophages in experimental pancreatic tumors**. R211 cells derived from spontaneous murine pancreatic tumors and genetically engineered to express Red-shifted luciferase reporter gene (R211-RSLuc) were engrafted in the pancreas of C57/BL6 mice as described in Material and Methods. At the end of the experiment, control mice and mice treated by 80µg of CL347 were killed, tumors were sampled and analyzed for CD3 and F4/80 (**A**). Quantification of CD3 (**B**) or F4/80 (**C**)-positive cells per field in control and mice treated by 80µg of CL347. Results are mean ± S.D. of ten fields from three different tumors. ***: p<0.005. Total RNA was extracted from control tumors or tumors treated by 80µg of CL347 and analyzed by Nanostring using the Ncounter murine inflammation panel as described in Material and Methods. Three tumors were used per group. **D**. Volcano plot of statistical significance (p< 10^-5^, 248 genes) against fold-change between CL347-treated and control mice. Grey dots: NS: not statistically significant; green dots: Log2 fold change, not statistically significant; red dots, Log2 fold change, not statistically significant. C2 and TNF were later excluded due to insufficient reads (<50) **E**. mRNA expression of selected genes with significant variations between control and CL347-treated tumors. Results are mean ± S.D. of three different tumors per group. *: p<0.05, **: p<0.01, ***: p<0.005.
