## Supplementary figures and images for "The antitumoral activity of TLR7 ligands is corrupted by the microenvironment of pancreatic tumors"

### supplemental file 1

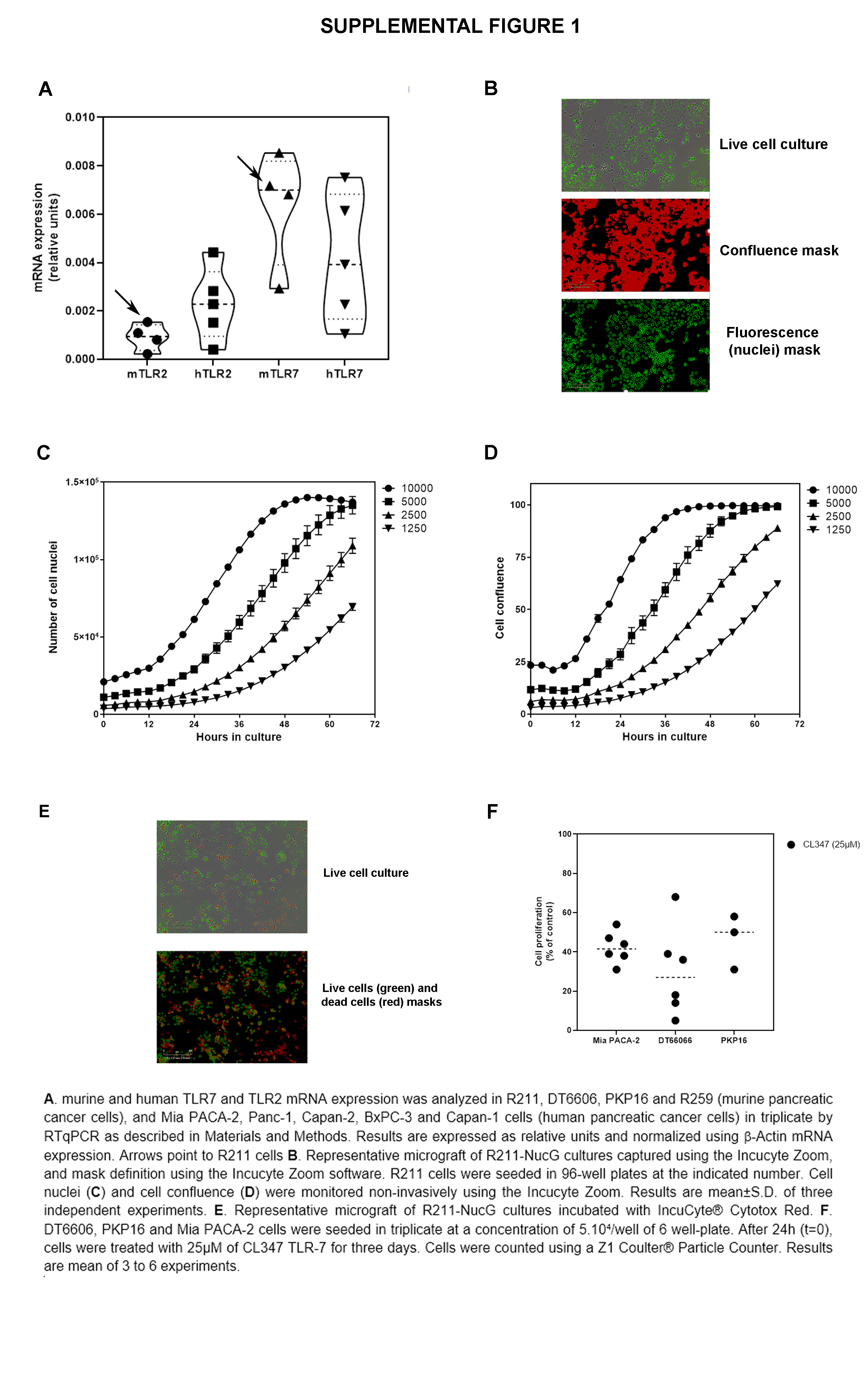
